# Influenza A Virus Interferes With Respiratory Syncytial Virus in Mice and Reconstituted Human Airway Epithelium

**DOI:** 10.1101/2025.02.19.639117

**Authors:** Shella Gilbert-Girard, Jocelyne Piret, Chantal Rhéaume, Julie Carbonneau, Nathalie Goyette, Christian Couture, Guy Boivin

## Abstract

Epidemiological studies suggest that respiratory syncytial virus (RSV) and influenza A virus (IAV) might interfere with each other. Viral interference mainly relies on interferon production elicited by a first virus that reduces the replication of a second virus. In this paper, we first investigated the interactions between RSV-A2 and influenza A(H1N1)pdm09 in BALB/c mice infected with each single virus or both viruses simultaneously or sequentially before, at the peak of interferon elicited by each virus or after that peak. IAV reduced by almost 3.0 logs the replication of RSV administered at the peak of interferon induced by influenza, but the opposite was not true. However, IAV-infected mice challenged with RSV or the vehicle lost more weight and had a lower survival rate compared to single infections. Interferon expression, cytokine levels and pulmonary inflammation were almost similar between groups. Disease worsening was attributed to an aggravation of IAV-induced pulmonary congestion following intranasal instillation of fluid (with or without RSV). In human airway epithelia, IAV also interfered with RSV replication. Viral interference was dependent on the timing and sequence of infections, but not on differential interferon susceptibilities. Overall, our results help to understand the mechanisms of interaction between two major respiratory viruses.

**Importance:** Respiratory syncytial virus and influenza virus may interfere with each other based on epidemiological studies. It is suggested that a first virus may induce the production of interferon and interfere with the replication of a second unrelated virus. Our data showed that influenza A virus interferes with respiratory syncytial virus replication in mouse lungs, but the opposite was not observed. In reconstituted human airway epithelia, viral interference was dependent on the timing and sequence of infections, but not on differential interferon susceptibilities. Understanding the mechanisms of interaction between respiratory viruses may help the development of prophylactic or therapeutic modalities.

## Introduction

Co-detections of two or multiple respiratory viruses are estimated to account for ≥10% of positive respiratory specimens from outpatients and hospitalized individuals, of which 84% are from children aged ≤5 years (1). Some studies have reported an association between coinfections and severe disease outcome (2–5) whereas others showed no significant difference in disease burden between single or multiple virus infections (6–8). During coinfections, viruses can interact with each other in the upper and/or lower respiratory tracts.

Epidemiological studies have shown that the likelihood of co-detecting antigenically distinct viruses in patient samples can be similar, lower or higher than that expected from random associations (9–16) suggesting neutral, negative (antagonistic) or positive (additive or synergistic) viral interactions. Mechanisms of negative viral interactions (i.e., a first viral infection decreases infection/replication of a second unrelated virus) could involve a competition between the two viruses for entry receptors and cellular resources based on *in vitro* or *in silico* studies (17, 18). During infection, host innate immune cells play an important role in the control of viral replication. It is thus more often suggested that negative viral interactions are mediated by the transient interferon (IFN) response elicited by the first virus through a mechanism known as viral interference (19). It is proposed that viruses that are strong inducers of an IFN response could reduce the infection/replication of viruses that show susceptibility to IFNs. The strength of the inhibitory effect is dependent on the kinetics of IFN production induced by the first virus, and thus on the time interval between the two viral infections. As respiratory viruses have developed their own immune evasion mechanisms to escape host IFN response (20–22), viral interference effects can be also dependent on the susceptibility of each single virus to IFN. Mechanisms of positive viral interactions (i.e., a first viral infection increases infection/replication of a second unrelated virus) are much less described for respiratory viruses (23). Viral interactions can be unidirectional or bidirectional, and thus depend on the sequence of infections.

Influenza and the respiratory syncytial virus (RSV), which are responsible for acute respiratory tract infections, are leading causes of morbidity and mortality in infants and in the elderly (24). A negative interaction between influenza A virus (IAV) and RSV is supported by the fact that the probability of detecting both viruses in swab specimens of patients was lower than expected from random association (10, 25). Furthermore, observational studies have reported that the emergence of the 2009 pandemic influenza A(H1N1) virus (A(H1N1)pdm09) delayed peaks of RSV seasonal activity in several countries in 2009-10 (26–31).

In this paper, we first investigated potential interference effects between influenza A(H1N1)pdm09 and RSV-A2 in a BALB/c mouse model. Our experimental approach was based on the kinetics of IFN response in mouse lungs, which markedly differed during RSV and IAV infections with peaks occurring on days 1 and 4 post-infection (p.i.), respectively. We thus evaluated viral interactions between RSV and IAV in mice infected intranasally with both viruses simultaneously or sequentially at time intervals based on the peak of IFN elicited by each virus. Interestingly, a prior infection with IAV reduced by almost 3.0 logs the RSV load when given at the peak of IFN induced by influenza. In contrast, a less than 0.5 log decrease in the replication of IAV was observed after a first RSV infection. However, IAV-infected mice challenged with RSV or the vehicle 1 day or 4 days later lost more weight and had a lower survival rate compared to single infections. No major differences in the IFN expression, cytokine protein levels and pulmonary inflammation were found between IAV-infected mice that were challenged with RSV compared to single RSV infection. The deterioration of the clinical outcome was associated with an aggravation of IAV-induced pulmonary congestion following intranasal instillation of fluid (but not to the second viral infection). We further confirmed the interference pattern between IAV and RSV in nasal and bronchiolar human airway epithelia (HAEs). As shown in mice, the interference of IAV on RSV in nasal HAEs was related to the kinetics of IFN induced by influenza, but not to differential susceptibility to type I and III IFNs. Overall, these results help to understand how two major respiratory viruses interact together and could guide future prophylactic and therapeutic modalities.

## Results

### The kinetics of IFN expression differs in lungs of mice infected with RSV and IAV

To highlight potential interference effects between RSV and IAV, we first determined the kinetics of type I and type III IFN expression by reverse transcription digital droplet polymerase chain reaction (RT-ddPCR) in the lungs of BALB/c mice infected intranasally with RSV-A2 or influenza A(H1N1)pdm09. During RSV infection, the expression of IFN-β and IFN-λ2/3 was higher than that of IFN-α, with both peaks occurring on day 1 p.i. (Fig. 1A). During IAV infection, IFN-β was the most expressed compared to IFN-α and IFN-λ2/3, with all peaks observed on day 4 p.i. (Fig. 1B). The expression of IFN-α and IFN-β in IAV-infected mice was higher than the basal levels (on day 0; Fig. 1B) between day 1 and day 4 p.i. and between day 2 and day 6 p.i., respectively. Thus, the kinetics of IFN expression induced by RSV and IAV in mouse lungs differs with peaks occurring on day 1 and day 4 p.i., respectively.

**FIG 1.**
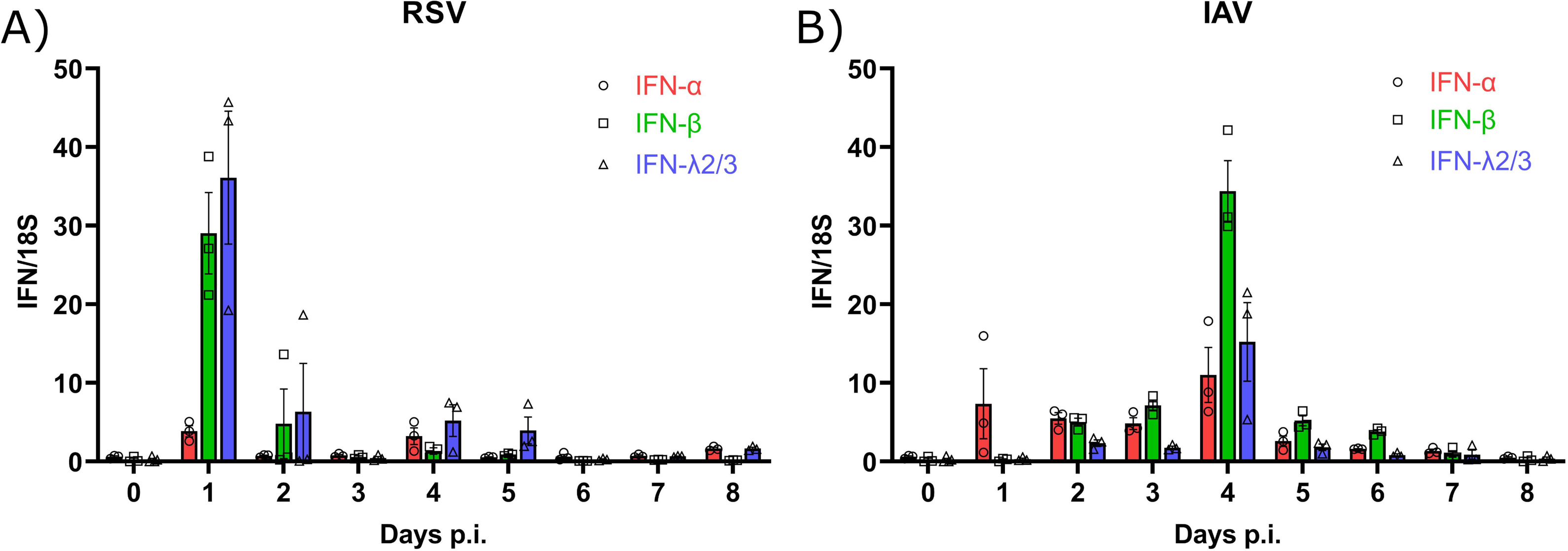
Kinetics of interferon (IFN) expression in the lungs of mice infected with respiratory syncytial virus (RSV) and influenza A virus (IAV). Interferon (IFN)-α and -β (type I) and IFN-λ2/3 (type III) expression was measured in lung homogenates by RT-ddPCR daily after RSV (A) and IAV (B) infection. Results are expressed as the mean of the ratio of IFN mRNAs over that of the 18S housekeeping gene (both in copies per µL) ± SEM of 3 mice per group from a single experiment.

### A prior infection with IAV reduces the replication of RSV in mouse lungs but the opposite is not true

We next investigated the interactions between RSV and IAV based on the kinetics of IFN expression induced by each virus in our mouse model. Mice were infected intranasally with each single virus or coinfected simultaneously or sequentially with both viruses (Fig. 2). For sequential infections, we selected time intervals corresponding to the peaks of IFN response induced by RSV (day 1 p.i.) and IAV (day 4 p.i.) in mouse lungs (Fig. 2). To evaluate interference effects between RSV and IAV, we determined the viral RNA loads of both viruses in lung homogenates by quantitative reverse transcription polymerase chain reaction (RT-qPCR). During simultaneous coinfections, the viral loads of RSV (Fig. 3A) and IAV (Fig. 3B) were similar to those of single infections. Interestingly, a prior IAV infection reduced by 0.9 log (with a 1-day interval; p<0.05) and almost 3.0 logs (with a 4-day interval; p<0.01) the viral load of RSV in mouse lungs compared to single RSV infection on day 4 after challenge (Fig. 3A). In contrast, a prior infection with RSV only slightly decreased the replication of IAV (by less than 0.5 log with a 4-day interval; p<0.01) compared to single IAV infection on day 4 after challenge (Fig. 3B). Thus, IAV interferes with the replication of RSV when the latter is administered at the peak of IFN expression induced by influenza.

**FIG 2.**
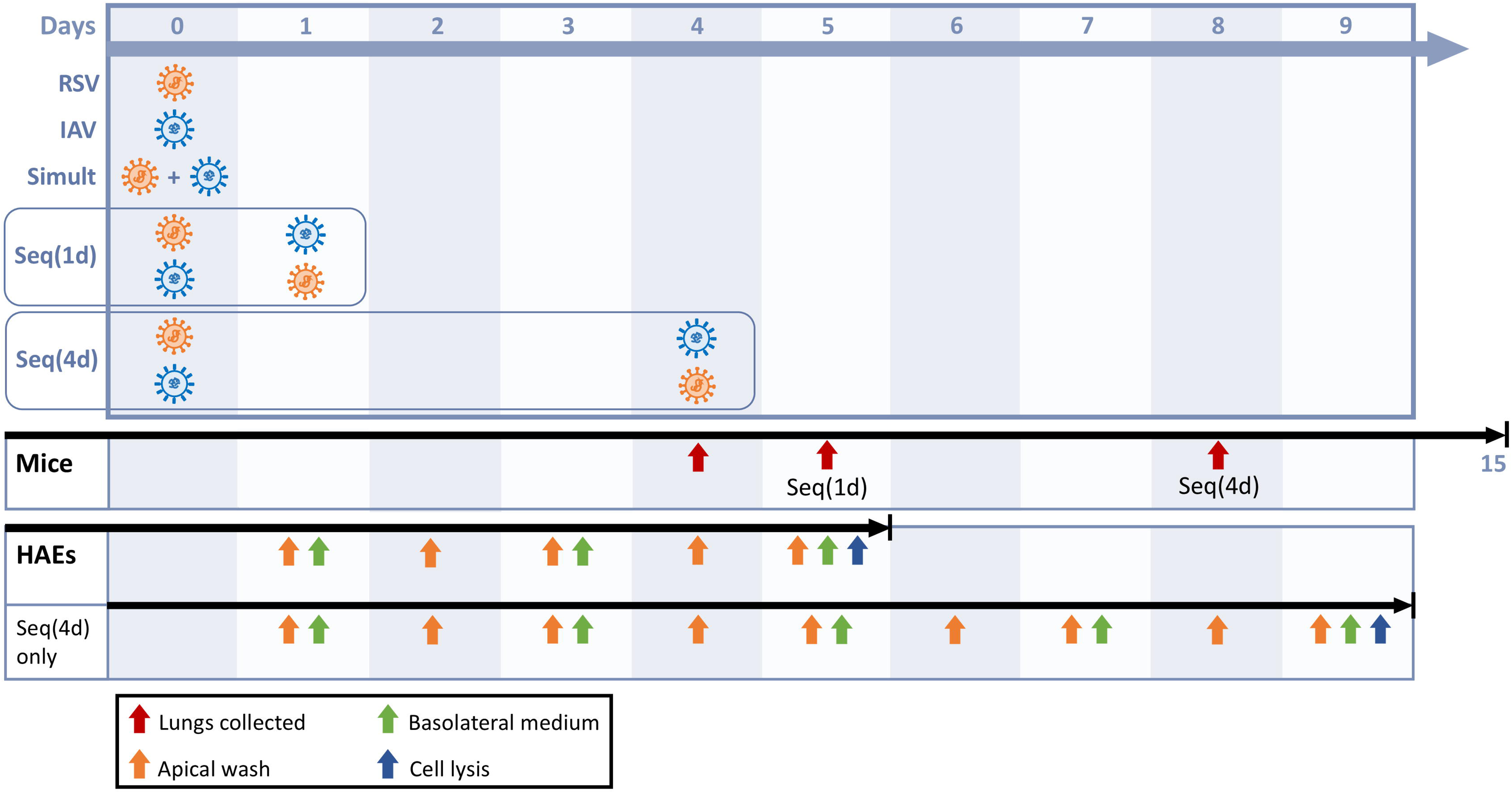
Timelines of single infections and coinfections with respiratory syncytial virus (RSV) and influenza A virus (IAV) in mice and human airway epithelia (HAEs). Mice and HAEs were infected with each single virus (RSV in orange and IAV in blue). In coinfection experiments, RSV and IAV were added either simultaneously (Simult) or sequentially with a 1-day (Seq(1d)) or a 4-day (Seq(4d)) interval. In mice, body weight changes and clinical signs of infection were monitored daily for 15 days. Lungs were collected 4 days after the last infection to determine viral RNA loads, interferon (IFN) expression, cytokine protein levels and histopathological evaluations. In HAEs, apical washes were collected daily (viral RNA loads), while basolateral medium was taken (IFN protein levels) and replaced by 500 µL of fresh media every 2 days. HAEs were lysed with RNA extraction buffer 5 days after the last infection, except for the Seq(1d) group (4 days after the second viral challenge) (interferon-stimulated gene expression).

**FIG 3.**
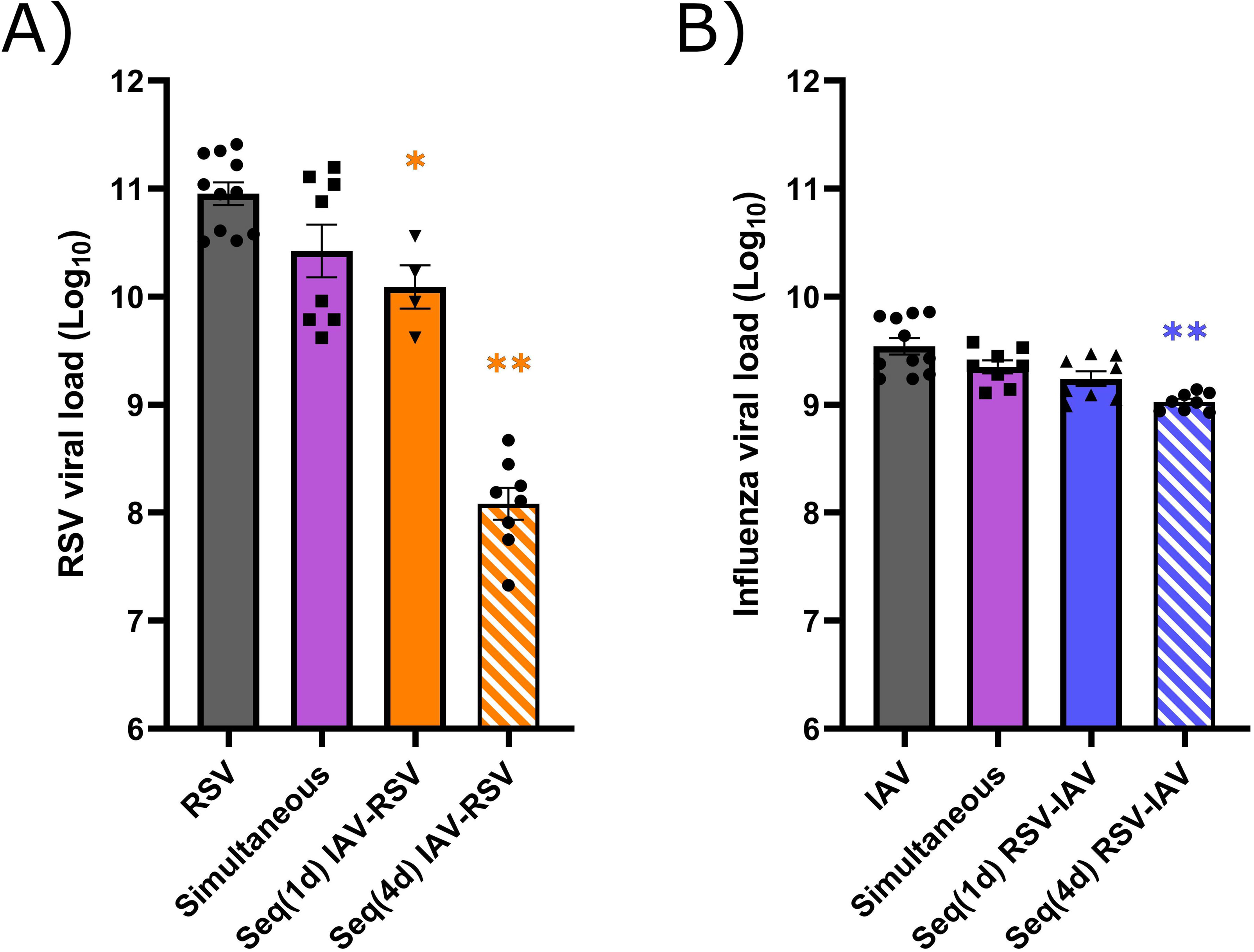
Viral RNA load in the lungs of mice infected with respiratory syncytial virus (RSV), influenza A virus (IAV) or both viruses. Mice were infected with RSV, IAV or both viruses simultaneously or sequentially with a 1-day (Seq(1d)) or a 4-day (Seq(4d)) interval. Viral RNA loads were determined in mouse lungs by RT-qPCR on day 4 after infection with RSV (A) or IAV (B). Results are expressed as the mean of the Log_10_ of total viral RNA copies in lung homogenates ± SEM of 4 to 11 mice per group from three independent experiments. * in orange, compared with RSV alone; * in blue, compared with IAV alone; *, p ≤ 0.05; **, p ≤ 0.01.

### Intranasal instillation of vehicle to IAV-infected mice worsens the clinical outcome

We also evaluated the clinical outcome of mice infected with RSV, IAV or coinfected simultaneously or sequentially with both viruses with a 1-day or a 4-day interval (Fig. 2). Mice that received the vehicle (minimal essential medium (MEM)) alone or sequential administrations of MEM, RSV or IAV followed by MEM with a 1-day or a 4-day interval were used as controls. Body weight changes and clinical signs of infection were monitored for 15 days after the first challenge. Maximum weight losses of mice infected with RSV and IAV alone occurred on day 6 and between days 7 and 8 p.i., respectively (Fig. 4A). The weight loss of mice coinfected simultaneously with the two viruses was more rapid and more severe compared to single infections (Fig. 4A). Mice infected sequentially with both viruses 1 day (Fig. 4B) or 4 days (Fig. 4C) later lost more weight compared to those infected with each single virus, especially when IAV was followed by RSV. The weight loss was more rapid in the Seq(1d) IAV-RSV group compared to the Seq(4d) IAV-RSV group. Notably, the weight loss pattern of mice that received intranasal administrations of IAV followed by MEM 1 day (Fig. 4B) or 4 days (Fig. 4C) later was almost similar to that seen during sequential infections with IAV and RSV at the same time interval. With a 1-day interval, RSV-infected mice that received the vehicle lost less weight than those infected with RSV followed by IAV whereas with a 4-day interval, the weight loss was comparable between both groups until day 6 p.i. and was lower in the Seq(4d) RSV-MEM group thereafter.

**FIG 4.**
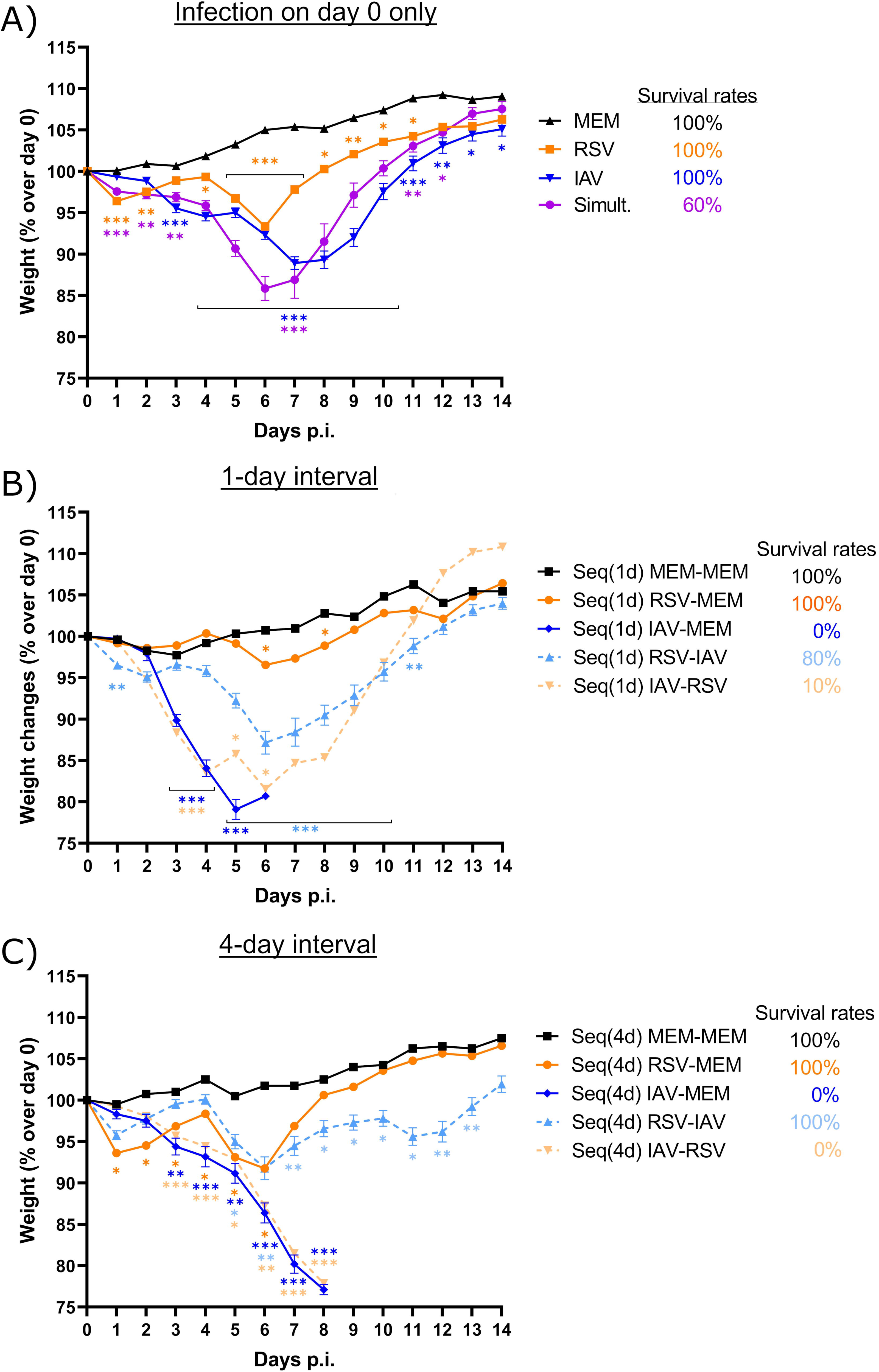
Body weight changes and survival rates of mice infected with respiratory syncytial virus (RSV), influenza A virus (IAV) or both viruses. Mice received intranasal administration of (A) the vehicle (minimum essential medium; MEM), RSV or IAV alone, or both viruses simultaneously on day 0; (B) the vehicle, RSV or IAV followed by the vehicle, or both viruses with a 1day interval (Seq(1d)); (C) the vehicle, RSV or IAV followed by the vehicle or both viruses with a 4-day interval (Seq(4d)). Body weight changes and survival rates were recorded daily from day 0 to day 14 post-infection (p.i.). Results are expressed as the mean of the percentage of the weight over that of day 0 ± SEM of 8-19 mice per group from three independent experiments. * in orange, compared with RSV alone; * in blue, compared with IAV alone; *, p ≤ 0.05; **, p ≤ 0.01; ***, p≤ 0.001.

All mice infected with IAV and RSV alone survived the infection (Fig. 4A). However, the survival rate of mice infected simultaneously with both viruses was reduced to 60% (Fig. 4A). During sequential infections with RSV followed by IAV, the survival rate decreased to 80% with a 1-day interval (Fig. 4B) whereas all mice survived the infection with a 4-day interval (Fig. 4C). The survival rates of mice infected with IAV followed by RSV 1 day and 4 days later were as low as 10% (Fig. 4B) and 0% (Fig. 4C), respectively. Importantly, all IAV-infected mice that received the vehicle 1 or 4 days later also succumbed to infection (Fig. 4, B and C) whereas anesthesia with isoflurane (without any intranasal instillation) did not increase mortality (data not shown). This indicates that the increased mortality seen in mice infected with IAV followed by RSV was not due to RSV infection by itself but rather to the vehicle that was administered by intranasal instillation.

### The reduced RSV load in IAV-infected mice is associated with a low IFN response

It is well acknowledged that the host IFN response is involved in the control of viral replication. However, viral infections or coinfections could also induce an exaggerated and prolonged IFN response, which could lead to a deterioration of the host clinical outcome (32). We thus determined the expression of IFN-α (Fig. 5A), IFN-β (Fig. 5B) and IFN–λ2/3 (Fig. 5C) in lung homogenates of the different mouse groups by RT-ddPCR on day 4 after the last infection. In single infections, IAV caused a higher type I IFN response than RSV (p<0.05 for IFN-β). Simultaneous coinfection with IAV and RSV induced an IFN response similar to both single infections, with some mice exhibiting higher expression levels especially for IFN-λ2/3. Compared to single infection with RSV, IFN-α and IFN-λ2/3 expression was decreased in the Seq(4d) IAV-RSV group on day 4 after RSV challenge. On day 4 post-IAV infection, IFN-α expression was lower in the Seq(1d) RSV-IAV group than in mice infected with influenza. Therefore, the reduced RSV load in the Seq(4d) IAV-RSV group is associated with lower IFN expression in the lungs.

**FIG 5.**
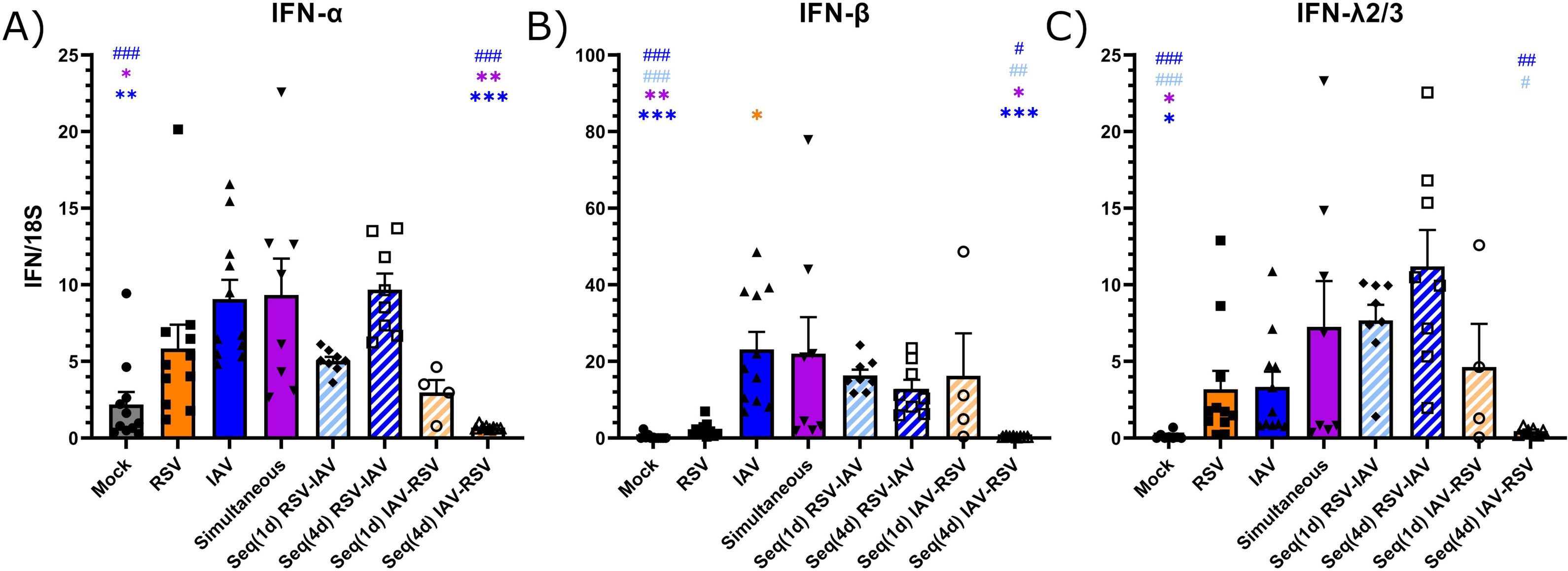
Expression of interferon (IFN) in the lungs of mice infected with respiratory syncytial virus (RSV), influenza A virus (IAV) or both viruses. Mice received intranasally the vehicle (mock), RSV or IAV, or both viruses simultaneously or sequentially with a 1day (Seq(1d)) or a 4-day (Seq(4d)) interval. IFN-α (A), IFN-β (B) and IFN-λ2/3 (C) mRNAs were determined in lung homogenates by RT-ddPCR on day 4 after the last infection. Results are expressed as the ratio of IFN mRNAs over that of the 18S housekeeping gene (both in copies per mL) ± SEM of 4-8 mice per group from three independent experiments. *, compared with single or simultaneous infection; ^#^, compared with sequential coinfection; *, ^#^, p ≤ 0.05; **, ^##^, p ≤ 0.01; ***, ^###^, p ≤ 0.001.

### The deterioration of the clinical outcome of IAV-RSV infected mice is not associated **with an increased cytokine production**

To evaluate whether the clinical outcome of mice assigned to the different coinfection conditions would be associated with changes in cytokine production compared to single infections, we measured the levels of IFN-γ, IL-1α, IL-1β, IL-6, IL-10 and TNF-α in lung homogenates by magnetic microbead immunoassays on day 4 after the last infection (Fig. 6). Single infections with RSV and IAV did not result in a markedly different cytokine production, except that RSV induced higher secretion of the anti-inflammatory cytokine IL-10 than IAV (p<0.05) and IAV showed a tendency to induce higher levels of the inflammatory cytokine IL-1β than RSV. Coinfections often led to similar or slightly higher cytokine levels than single infections, except that the production of IFN-γ and IL-10 was significantly higher in the Seq(1d) RSV-IAV group than in the IAV group (p<0.001 for both) on day 4 post-IAV challenge. Thus, the deterioration of the clinical outcome of mice infected with IAV followed by RSV is not associated with major changes in cytokine production in the lungs.

**FIG 6.**
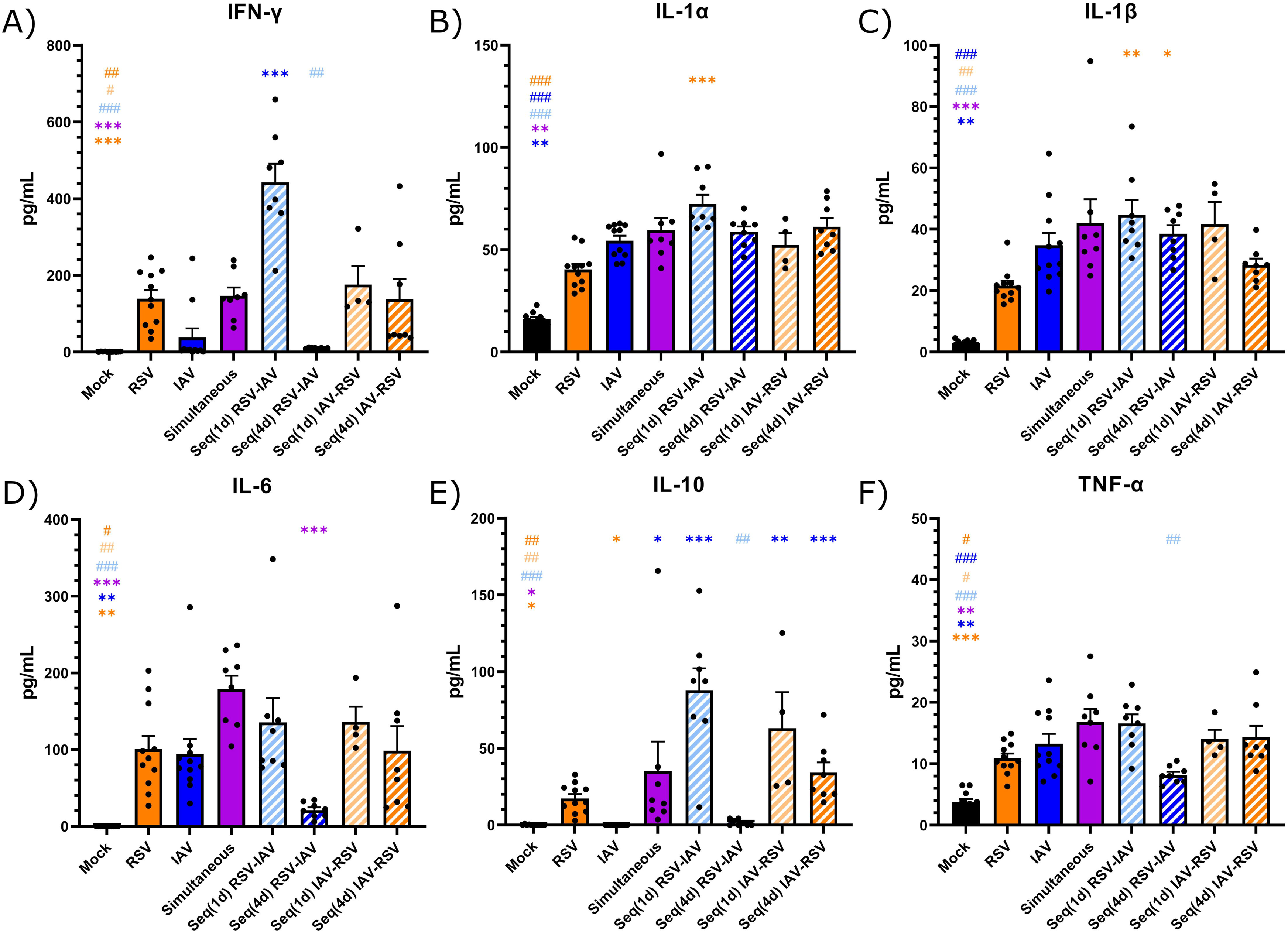
Production of cytokines in the lungs of mice infected with respiratory syncytial virus (RSV), influenza A virus (IAV) or both viruses. Mice received intranasally the vehicle (mock), RSV or IAV, or both viruses simultaneously or sequentially with a 1-day (Seq(1d)) or a 4-day (Seq(4d)) interval. Production of IFN-γ (A), IL-1α (B), IL-1β (C), IL-6 (D), IL-10 (E), TNF-α (F) was determined in lung homogenates by magnetic microbead immunoassays on day 4 after the last infection. Results are expressed as the mean of protein concentration in pg/mL ± SEM of 4-11 mice per group from three independent experiments. *, compared with single or simultaneous infection; ^#^, compared with sequential coinfection; *, ^#^, p ≤ 0.05; **, ^##^, p ≤ 0.01; ***, ^###^, p ≤ 0.001.

### The deterioration of the clinical outcome in IAV-RSV infected mice is not associated with an increased pulmonary inflammation

We then evaluated whether the clinical outcome of mice assigned to the different coinfection conditions would be associated with changes in pulmonary damage and inflammation compared to single infections. Fixed lungs were harvested 4 days after the last infection and processed for histopathological evaluation. The mean scores for inflammation in lung tissues of the different mouse groups are shown in Table 1. The mean total inflammation score was higher for mice infected with RSV than those infected with IAV (3.9 *versus* 2.5). However, histopathological evaluation of lung tissues did not reveal important changes in bronchial/endobronchial, peribronchial, perivascular, interstitial, pleural and intra-alveolar inflammation intensity as well as in total inflammation score between mice infected with one or two viruses. Thus, the deterioration of the clinical outcome of mice infected with IAV followed by RSV is not associated with a major inflammatory response in lung tissues.

**Table 1.** Histopathological evaluation of lung tissues of mice infected with respiratory syncytial virus, influenza A virus or both viruses on day 4 after the last infection

| Groups | Bronchial/<br>endobronchial | Peribronchial | Perivascular | Interstitial | Pleural | Intra-alveolar | Total inflammation<br>score |
| --- | --- | --- | --- | --- | --- | --- | --- |
| Mock | 0.0 | 0.0 | 0.0 | 0.0 | 0.0 | 0.0 | 0.0 |
| RSV | 0.3 | 0.4 | 0.9 | 1.1 | 0.4 | 0.9 | 3.9 |
| IAV | 0.8 | 0.9 | 0.6 | 0.1 | 0.0 | 0.1 | 2.5 |
| Simult. | 1.0 | 1.0 | 0.4 | 0.8 | 0.3 | 0.8 | 4.1 |
| Seq(1d) RSV-IAV | 0.6 | 1.0 | 0.6 | 0.9 | 0.0 | 0.9 | 4.0 |
| Seq(4d) IAV-RSV | 1.0 | 0.8 | 0.7 | 0.5 | 0.0 | 0.5 | 3.5 |

RSV, respiratory syncytial virus; IAV, influenza A virus; Mock, mice that received an intranasal instillation of the vehicle; Simult., simultaneous coinfection; Seq(1d), sequential infection with 1-day interval; Seq(4d), sequential infection with 4-day interval. Scores are the mean of 3-4 mice per group from a single experiment.

### An intranasal instillation of fluid aggravates IAV-induced pulmonary congestion

As an intranasal instillation of vehicle was sufficient to deteriorate the clinical outcome of IAV-infected mice, we evaluated the development of pulmonary congestion in mice infected with IAV and RSV by determining the lung wet-to-dry weight ratio on days 1, 4, 7 and 14 p.i. In mice infected with RSV, this ratio was higher on day 1 p.i. and decreased thereafter suggesting that the formation of pulmonary congestion is rapid and transient (Fig. S1). Some RSV-infected mice challenged with IAV 1 day later (20%), which corresponds to the peak of pulmonary congestion, showed a deterioration of the clinical outcome, but not with a 4-day interval (after resolution of pulmonary congestion). In mice infected with IAV, the lung wet-to-dry weight ratio increased steadily from day 1 to day 7 p.i. indicating that pulmonary congestion develops more slowly but steadily compared to RSV (Fig. S1). We may thus suggest that the intranasal instillation of vehicle (with or without RSV) to IAV-infected mice may have further aggravated the pulmonary congestion leading to disease worsening. Pulmonary congestion in IAV-infected mice resolved by day 14 p.i. (Fig. S1).

### IAV interferes with the replication of RSV in nasal and bronchiolar HAEs

We then used *ex vivo* models of upper and lower respiratory tracts based on reconstituted nasal and bronchiolar HAEs to further study the interactions between RSV-A2 and influenza A(H1N1)pdm09. These HAEs are composed of fully differentiated human cell types such as ciliated, goblet, basal and club (bronchiolar only) cells. Despite the fact that they do not contain immune cells, these epithelia can secrete cytokines and other immune mediators, and allow to study the early steps of viral interactions. Both viruses infected successfully nasal and bronchiolar HAEs and their kinetics of replication followed similar trends (shown in Fig. 7 and Fig. S2, respectively).

**FIG 7.**
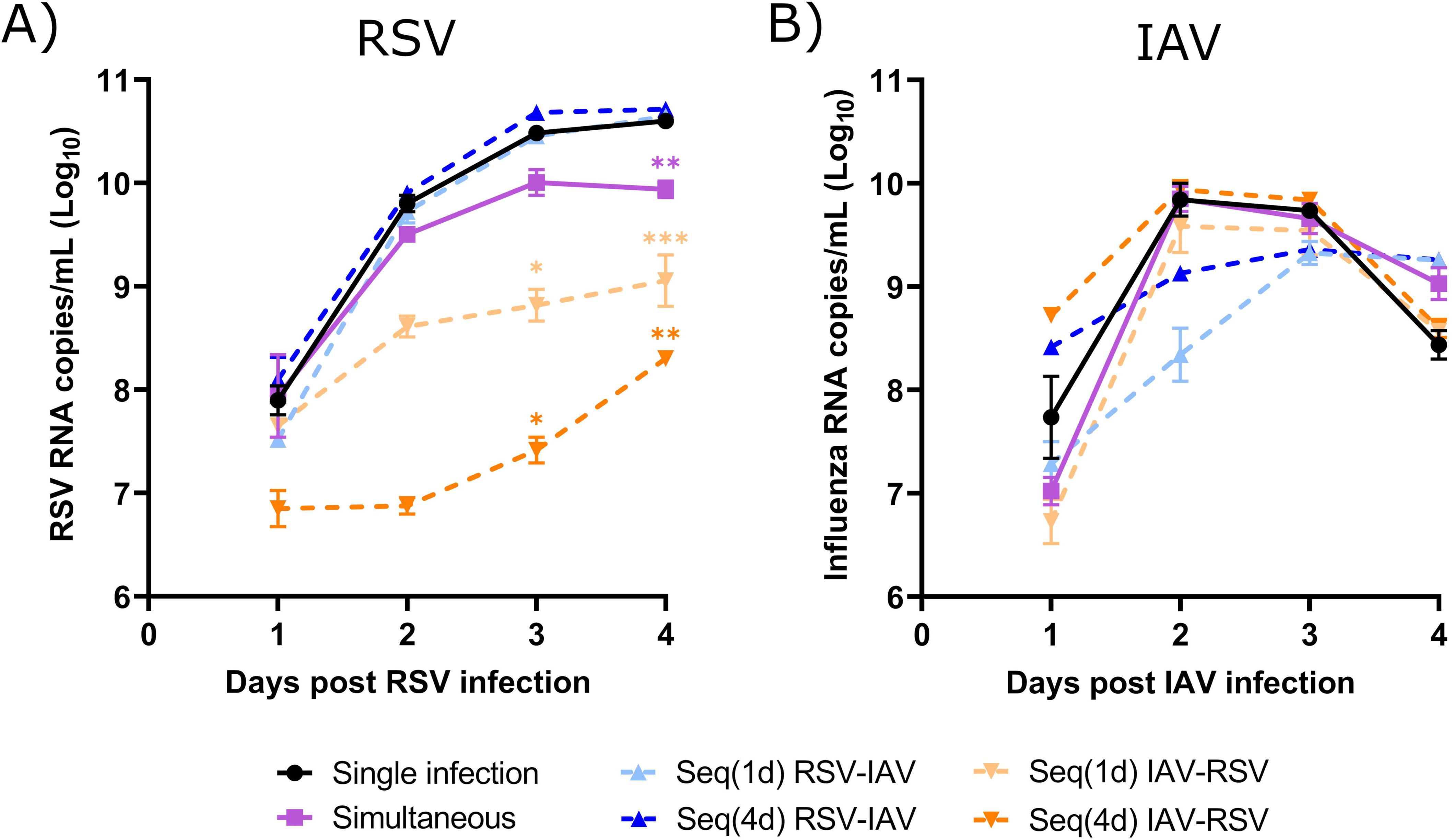
Viral interference between respiratory syncytial virus (RSV) and influenza A virus (IAV) in nasal human airway epithelia (HAEs). Nasal HAEs were infected with RSV, IAV, or both viruses simultaneously or sequentially with a 1-day (Seq(1d)) or a 4-day (Seq(4d)) interval. Viral RNA loads were determined in apical washes by RT-qPCR on a daily basis. Days post-infection (p.i.) represent the time after infection with either RSV (A) or IAV (B). Results are expressed as the mean of the Log_10_ of viral RNA copies per mL ± SEM of 3-5 nasal HAE inserts from two independent experiments. *, p ≤ 0.05; **, p ≤ 0.01; ***, p ≤ 0.001 compared to single infection.

Nasal HAEs were infected with each single virus or coinfected with both viruses simultaneously or sequentially 1 day or 4 days apart (Fig. 2). The viral load of IAV reached its peak on day 2 p.i., whereas that of RSV increased more slowly to reach higher levels and possibly a plateau at later times after infection. When nasal HAEs were infected simultaneously with both viruses, the viral RNA load of RSV was only reduced by 0.7 log (p<0.01) compared to single infection on day 4 p.i. (Fig. 7A), whereas the replication of IAV was not significantly affected (Fig. 7B). Compared to single infection, the viral load of RSV in IAV-infected nasal HAEs was markedly decreased by 1.7 log (p<0.05; 1-day interval) and 3.0 logs (p<0.05; 4-day interval) on day 3 p.i. and by 1.5 log (p<0.001; 1-day interval) and 2.3 logs (p<0.01; 4-day interval) on day 4 p.i. (Fig. 7A). In contrast, a prior infection of nasal HAEs with RSV showed only a tendency to reduce the viral load of IAV with a 1-day time interval (Fig. 7B).

The viral interference effects between IAV and RSV was confirmed in bronchiolar HAEs. A prior infection of bronchiolar HAEs with IAV also markedly decreased the replication of RSV compared to single infection (Fig. S2A), especially with a 1-day interval (over 4.0 logs of reduction between day 2 and day 4 p.i. (p<0.01)). In bronchiolar HAEs infected with IAV followed by RSV 4 days later, the viral load of RSV was significantly reduced by 1.4 logs (p<0.05) and 3.1 logs (p<0.01) on days 1 and 2 p.i., respectively, but increased steadily thereafter. In contrast, the replication of IAV in bronchiolar HAEs was not affected by a prior challenge with RSV (Fig. S2B). Thus, a prior infection of nasal and bronchiolar HAEs with IAV reduces the replication of RSV.

### The kinetics of IFN production differ in nasal HAEs infected with RSV and IAV

We next evaluated whether the viral interference of IAV on RSV would be related to the kinetics of IFN production induced by influenza. The production of IFN-β, IFN-λ1 and IFN-λ2 proteins was measured in the basolateral medium of nasal HAEs infected with each single virus by magnetic microbead immunoassays on a daily basis for 5 days. Infection of nasal HAEs with IAV resulted in a faster production of IFN-β (Fig. 8A), IFN-λ1 (Fig. 8B) and IFN-λ2 (Fig. 8C) compared to RSV. Peaks of IFN levels elicited by IAV occurred between days 2 and 3 p.i. and were followed by a quick drop thereafter. In contrast, the IFN response induced by RSV reached a plateau with persistent levels between days 3 to 5 p.i. As shown in mouse studies, the viral interference of IAV on the replication of RSV is related to the kinetics of IFN production induced by influenza in nasal HAEs.

**FIG 8.**
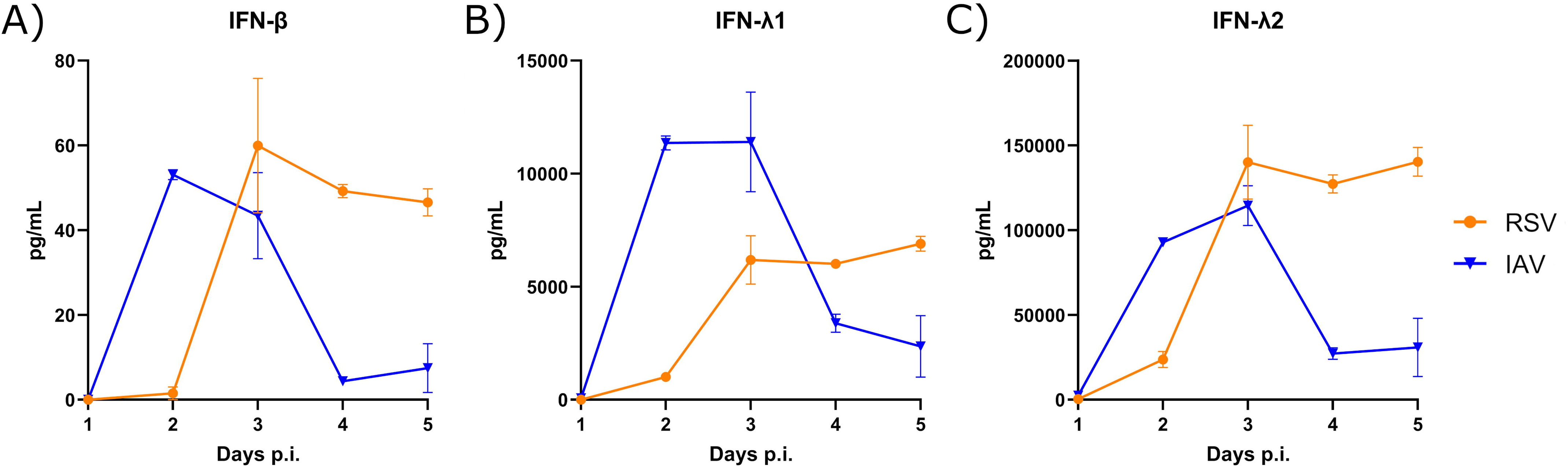
Interferon (IFN) protein production in nasal human airway epithelia (HAEs) infected with respiratory syncytial virus (RSV) and influenza A virus (IAV). IFN-β (A), IFN-λ1 (B) and IFN-λ2 (C) protein production was measured in basolateral medium by magnetic microbead immunoassays daily for 5 days post-infection (p.i.). Results are expressed as the mean amount of IFN proteins in pg per mL ± SEM of 2-4 nasal HAE inserts from two independent experiments.

### The reduced RSV load in IAV-infected nasal HAEs is associated with a low IFN response

We then determined IFN-β, IFN-λ1 and IFN-λ2 protein levels in the basolateral medium of nasal HAEs infected with each single virus or coinfected with the two viruses simultaneously or sequentially 1 day or 4 days apart (Fig. 2). All measurements were done on day 5 after the last infection, except for the Seq(1d) groups (day 4 after the second viral challenge). No IFN was detected in uninfected HAEs (data not shown). Single infections of nasal HAEs with IAV and RSV resulted in a lower IFN-β response (Fig. S3A) than that of IFN-λ1 (Fig. S3B) and IFN-λ2 (Fig. S3C). RSV showed a tendency to cause a higher production of all three IFNs compared to IAV. The IFN response induced during simultaneous coinfections with RSV and IAV was almost similar to that of RSV alone. Sequential infection of HAEs with RSV followed by IAV 1 day later induced a stronger IFN production than single infection with IAV (p<0.05 for IFN-β and IFN-λ2). Sequential infections of HAEs with IAV followed by RSV at both time intervals resulted in a lower IFN production than that induced by RSV alone, especially for the Seq(4d) IAV-RSV group. Thus, the reduced RSV load in IAV-infected HAEs is associated with lower levels of IFN proteins.

### Interferon-stimulated gene expression follows the IFN response in HAEs coinfected with IAV followed by RSV

We then evaluated the expression of four interferon-stimulated genes (ISGs) acting on different steps of viral infection (i.e., OAS1, IFITM3, ISG15 and MxA) in lysates of nasal HAEs infected with each single virus or coinfected with the two viruses simultaneously or sequentially 1 day or 4 days apart (Fig. 2). All determinations were done by RT-ddPCR on day 5 after the last infection, except for the Seq(1d) groups (day 4 after the second viral challenge). RSV showed a tendency to induce a higher ISG expression than IAV, especially for OAS1, ISG15 and MxA (Fig. 9). The expression of ISGs was almost similar during simultaneous coinfection than during single infection with RSV. The expression of IFITM3, ISG15 and MxA was higher in HAEs infected with RSV followed by IAV 1 day apart than with IAV alone (but determinations were made on day 4 and day 5 post-IAV, respectively); this increase was significant for ISG15 (p<0.001). The expression of all 4 ISGs was decreased in HAEs infected sequentially with IAV followed by RSV 4 days later compared to RSV alone, with a significant reduction of 0.6-and 0.7-fold for OAS1 and ISG15, respectively (p<0.001 for both). Therefore, the expression of ISGs in nasal HAEs infected with IAV followed by RSV was lower than that induced by RSV alone and follows almost the same trend as IFN protein production.

**FIG 9.**
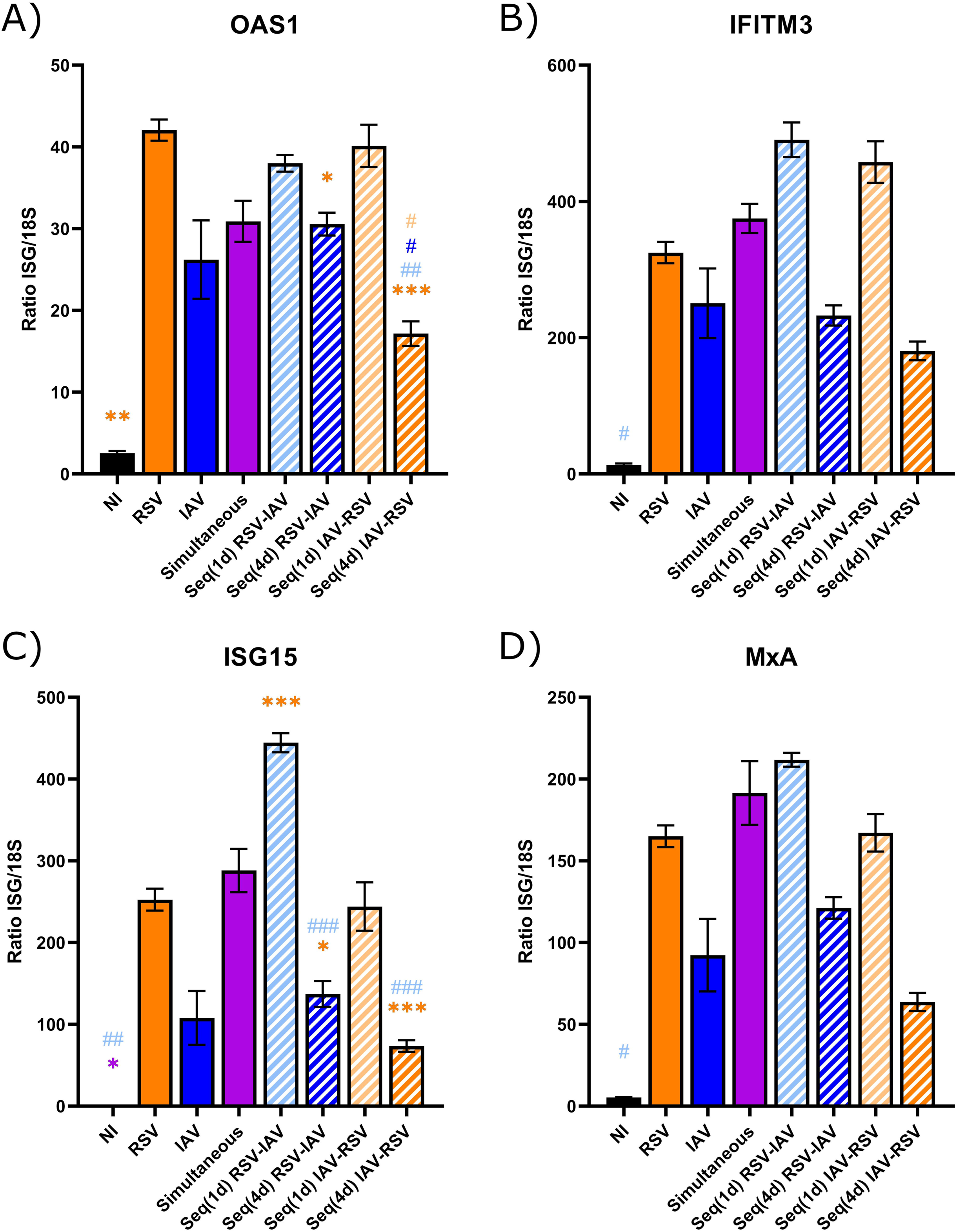
Interferon-stimulated gene (ISG) expression in nasal human airway epithelia (HAEs) infected with respiratory syncytial virus (RSV), influenza A virus (IAV) or both viruses. Nasal HAEs were infected with RSV, IAV, or both viruses simultaneously or sequentially with a 1-day (Seq(1d)) or a 4 day (Seq(4d)) interval. Non-infected (NI) HAEs were used in parallel. Expression of four ISGs (OAS1 (A), IFITM3 (B), ISG15 (C) and MxA (D)) was measured in cell lysates by RT-ddPCR on day 5 after the last infection, except for the Seq(1d) groups (4 days after the second viral challenge). Results are expressed as the mean of the ratio of ISG mRNAs over that of 18S housekeeping gene (both in copies per µL) ± SEM of 3 to 5 nasal HAE inserts from two independent experiments. *, compared with single or simultaneous infection; ^#^, compared with sequential coinfection; *, ^#^, p ≤ 0.05; **, ^##^, p ≤ 0.01; ***, ^###^, p ≤ 0.001.

### RSV and IAV have almost similar susceptibility to exogenous interferons

We then assessed the susceptibility of RSV and IAV to exogenous IFN-α2a, IFN-β and IFN-λ2 proteins in nasal HAEs. The viral RNA loads of RSV (Fig. 10A) and IAV (Fig. 10B) were markedly decreased in the presence of IFN-α2a and IFN-β. The effect of IFN-λ was less pronounced on both viral infections compared to those of type I IFNs, especially against RSV. Thus, the viral interference of IAV on the replication of RSV does not appear to be related to a difference in their susceptibility to IFN.

**FIG 10.**
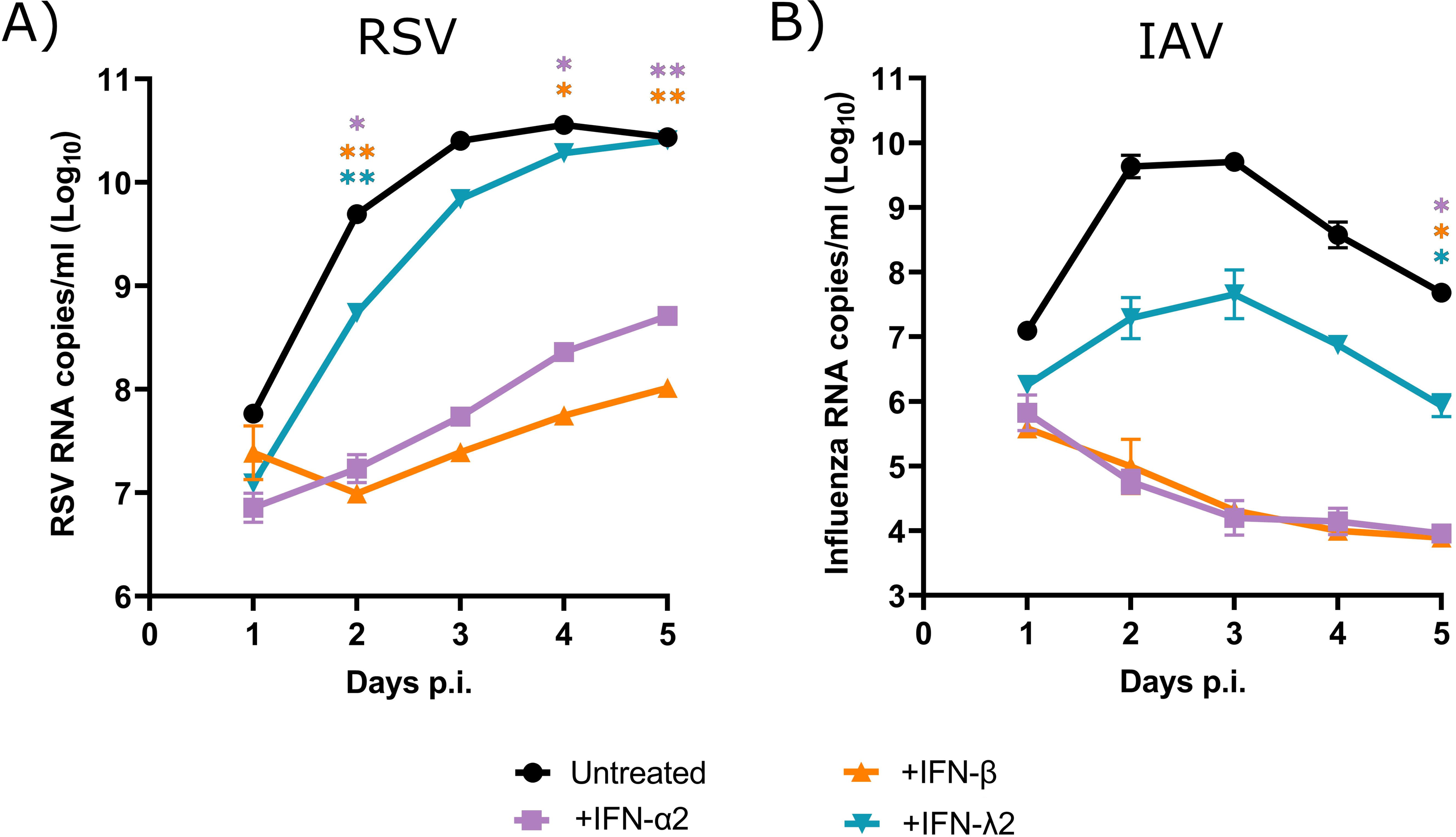
Susceptibility of respiratory syncytial virus (RSV) and influenza A virus (IAV) to recombinant interferon (IFN) proteins in nasal human airway epithelia (HAEs). Nasal HAEs were infected with RSV and IAV, in presence of IFN-α2a, IFN-β or IFN-λ2. Infected and untreated HAEs were used in parallel. The viral RNA loads of RSV (A) and IAV (B) were determined in apical washes by RT-qPCR daily for 5 days post-infection (p.i.). Results are expressed as the mean of the Log_10_ of viral RNA copies per mL ± SEM of 3 nasal HAE inserts in one experiment. A value of 60 copies/mL, corresponding to the detection limit of the assays, was attributed to samples with undetectable RNA levels. *, p ≤ 0.05; **, p ≤ 0.01 compared to untreated HAEs.

## Discussion

Viral interference effects between IAV and RSV have been suggested in epidemiological studies. In this paper, we investigated potential interference effects between RSV-A2 and influenza A(H1N1)pdm09 in BALB/c mice and HAEs to better understand their dynamics in the setting of coinfections.

To this end, we developed a mouse model of coinfection based on the kinetics of IFN expression elicited by the first virus. A prior infection with IAV significantly reduced the lung RSV load compared to a single infection. This reduction was higher when s mice were challenged with RSV at the peak of IFN expression induced by IAV (day 4 p.i.) rather than before that peak (day 1 p.i.). Thus, these data demonstrate that IAV interferes with the replication of RSV in mouse lungs and highlight the importance of the interval between the two infections, which is related to IFN expression elicited by the first virus. Such an interference effect of IAV on the replication of RSV according to the timing of infections has been previously reported in mice and ferrets (33–36).

In contrast, a prior infection of mice with RSV only slightly decreased the replication of IAV in mouse lungs with time intervals between the two infections corresponding to the peak of IFN expression elicited by RSV (day 1 p.i.) or after that peak (day 4 p.i.). Several studies reported mitigated results for the effects of RSV on the replication of IAV when evaluated in BALB/c mice. One study showed that the IAV load was reduced in the lungs of mice infected sequentially with RSV-A2 followed by A(H1N1) strain PR/8/34 with a 4-day interval (33) whereas another study did not report such an effect during sequential infections with the same RSV and IAV strains 1 day apart (34). In a third study, coinfection of mice with RSV-A2 and A/H3N2/Perth/16/09 with a 1-day interval resulted in lower lung pathology compared to influenza alone, as assessed by histopathogical evaluation (36). Differences in viral strains, viral inoculum, timing between the two infections and age/body weight of BALB/c mice could be responsible for such discrepancies. Nevertheless, in our mouse model, infection with RSV followed by IAV 4 days later, which resulted in a slight reduction of IAV replication in the lungs, was also associated with lower body weight loss at the peak of infection (between days 6 to 8 p.i.) compared with single IAV infection. A prior infection with RSV that only slightly reduces the replication of IAV in the lungs could thus attenuate the disease outcome compared to single influenza infection as shown by others (33–35).

However, all mice infected with IAV followed by RSV lost more weight and reached the limit point requiring their euthanasia, which is in sharp contrast with animals infected with each single virus that all survived the infection. Other groups also reported that sequential infection of mice with IAV followed by RSV deteriorates the disease outcome compared to single infections (34). However, the deterioration of the clinical outcome of mice infected with IAV followed by RSV could not be explained by an uncontrolled and prolonged IFN expression, changes in cytokine protein production or increased inflammation scores in lung tissues compared to infection with RSV alone.

Influenza virus, and to a lesser extent RSV, were shown to inhibit alveolar fluid clearance in the lungs of BALB/c mice (37). The mechanisms involved a reduction of the functions of the epithelial sodium channel and the cystic fibrosis transmembrane regulator in murine respiratory and alveolar epithelia (38). The determination of the lung wet-to-dry weight ratio showed that RSV infection caused a pulmonary congestion that rapidly resolved. In contrast, IAV infection induced a long-lasting and steadily increasing pulmonary congestion, which may explain that intranasal instillation of an additional volume of liquid (vehicle with or without RSV) to IAV-infected mice may lead to a worse outcome. Our hypothesis is that the second intranasal instillation of fluid increased the pulmonary congestion induced by IAV resulting in acute respiratory distress syndrome and a worse disease outcome. Furthermore, RSV-infected mice challenged with IAV 1 day later (at the peak of pulmonary congestion) showed a survival rate of 80% whereas all RSV-infected mice challenged with IAV 4 days later (after resolution of pulmonary congestion) survived infection. These results also suggest a relationship between the extent of pulmonary congestion induced by RSV and the deterioration of the clinical outcome following the second intranasal instillation of MEM (with or without IAV).

We then used *ex vivo* models based on reconstituted nasal and bronchiolar HAEs to further investigate the interactions between RSV and IAV during simultaneous or sequential infections 1 day or 4 days apart. We confirmed that IAV interfered with the replication of RSV in nasal HAEs. Also, similar results were observed in bronchiolar HAEs although a 1-day interval between IAV and RSV infections resulted in a stronger interference effect than a 4-day interval. An interference of influenza virus on the replication of severe acute respiratory syndrome coronavirus 2 (SARS-CoV-2) in HAEs has already been described in several reports (39–44). Our results also showed that RSV only slightly reduced the replication of IAV in nasal, but not in bronchiolar HAEs, suggesting differences in defense mechanisms induced by upper and lower respiratory epithelia. An interference of RSV on human metapneumovirus (45) and SARS-CoV-2 (39, 44) in nasal HAEs was nevertheless reported in other studies.

The interference effects of IAV on SARS-CoV-2 in HAEs have been partly attributed to the IFN response induced by the first virus (39–41, 44). We thus determined the kinetics of IFN production in nasal HAEs infected with RSV and IAV alone. We observed that the kinetics of IFN response induced by each virus correlated with their kinetics of replication. Indeed, IAV, that reaches its peak of replication on day 2 p.i. followed by a marked decrease of viral load thereafter, elicited a fast and short IFN response. This could lead to a more rapid establishment of an antiviral state in infected and uninfected surrounding cells, which may contribute to the ability of IAV to interfere with RSV. In contrast, RSV, that shows a steady replication between days 2 and 4, induced a sustained IFN production. However, this immune response could be elicited too late to reduce the replication of influenza even when the second viral challenge was done 4 days later. Our hypothesis is that the first infection with IAV triggers an immune response that prevents the replication of RSV, which cannot induce a proper IFN response. In this respect, the timing for harvesting lungs and basolateral medium is important to consider. In mice, IFN were quantified in lungs collected 4 days after RSV inoculation, which means 8 days after IAV infection. At this timepoint, IFN production elicited by IAV may have dropped and the response induced by RSV, which is not long-lasting, may be inhibited by viral interference due to IAV. The same phenomenon should occur in HAEs where IFNs were quantified in basolateral medium 5 days after adding RSV in the seq(4d) IAV-RSV group. In addition to the IFN response, alternative/additional mechanisms that may explain viral interference effects could involve competition for resources or cellular damage. We observed that IAV has a faster replication cycle than RSV in HAEs and it is possible that a faster IAV depletes resource availability for RSV in this *ex vivo* model. However, IAV does not seem to replicate faster than RSV in mouse lungs. This difference may be related to the fact that HAEs do not contain innate immune cells, which are key players in the control of viral replication.

As previously shown by our group (40), infection of nasal HAEs with IAV, as well as with RSV, resulted in a lower IFN-β response compared to those of IFN-λ1 and IFN-λ2. This is in accordance with previous reports showing that type III IFN is the main antiviral response in airway epithelium (46, 47). Consistent with its sustained IFN response, RSV showed a tendency to induce a more elevated production of all three IFNs with a slightly higher ISG expression than IAV on day 5 after infection. As shown in our mouse model, the reduced RSV load in HAEs infected sequentially with IAV and RSV 4 days later was associated with modest levels of IFN proteins. The expression of ISGs in HAEs was also low in this coinfection condition, especially for OAS1 and ISG15. Finally, no difference was found in the susceptibility of RSV and IAV to exogenous IFNs but the doses used in our study were higher than physiological concentrations, which may have prevented to observe a difference. Overall, these data suggest that the timing of IFN response induced by each virus and the sequence of viral infections are important parameters in the occurrence of viral interference effects rather than a difference in their susceptibility to IFNs.

The strength of our study is that we investigated the interference effects between RSV-A2 and influenza A(H1N1)pdm09 virus based on the kinetics of IFN response elicited by the first virus. We used both a mouse model (with innate and adaptive immune systems) and reconstituted nasal and bronchiolar airway epithelia of human origin. To confirm viral interference, we further studied the interactions between RSV and IAV during simultaneous and sequential infections before, at the peak of IFN response elicited by the first virus or after that peak. Nonetheless, our study has some limitations. The intranasal instillation of 50 µL of vehicle with or without RSV to IAV-infected mice might have resulted in an aggravation of the pulmonary congestion leading to a rapid deterioration of the clinical outcome. This suggests that an evaluation of pulmonary congestion in mice infected with each single virus (for example, as determined by the lung wet-to-dry weight ratio) is required when developing models to study the interactions between respiratory viruses during sequential overlapping infections. The volume used for the second viral challenge should be kept as small as possible to prevent an aggravation of the pulmonary congestion, if any. Finally, appropriate controls need to be included to assess whether intranasal instillation of the vehicle to mice infected with a first virus does not deteriorate the clinical outcome. In our experiments, mice were mock-infected with the vehicle (MEM) but culture medium of uninfected cells would have been more appropriate. Nevertheless, the concentrations of IFN or other cellular proteins that may have been present in our viral stocks have been markedly reduced by the dilutions required to get the appropriate inoculums for infection of mice or HAEs.

In conclusion, we showed that A(H1N1)pdm09 interferes with RSV-A2 in mouse and HAE models, whereas RSV has only a modest or no effect on influenza replication. The viral interactions between IAV and RSV are mainly related to the kinetics of IFN response elicited by each virus and the sequence of infections rather than to their susceptibility to type I and III IFNs. Other studies have already shown the major role of the IFN response in viral interference effects between several other pairs of viruses such as influenza and SARS-CoV-2 (39–44), rhinovirus and SARS-CoV-2 (41, 48–50), rhinovirus and influenza (51, 52) as well as RSV and human metapneumovirus (45). We cannot exclude, however, that viral interactions could be mediated by other mechanisms specific to each virus (53). A better understanding of the interactions between respiratory viruses could help in the prediction of epidemic peaks and pandemic waves through the improvement of mathematical models of viral transmission. Furthermore, this knowledge could promote the development of preventive and therapeutic options against respiratory viral infections based on the activation of innate immune response such as defective interfering particles of IAV (54).

## Materials and methods

### Cells and viruses

ST6Gal I Madin-Darby Canine Kidney cells overexpressing the 112-6 sialic acid receptor (MDCK 112-6) were obtained from Dr. Y. Kawaoka (University of Wisconsin, Madison, WI, USA) (55). MDCK 112-6 cells and HEp-2 (human epithelial carcinoma; CCL-23; American Type Culture Collection (ATCC), Manassas, VA, USA) cells were cultured in MEM (Invitrogen, Carlsbad, CA, USA) supplemented with 10% fetal bovine serum (FBS; Invitrogen) and 1% HEPES. Puromycin (7.5 µg/mL) was also added to the culture medium of MDCK 112-6 cells. Nasal (MucilAir™, pool of donors, EP02MP) and bronchiolar (SmallAir™, single donor, EP21SA) reconstituted HAE inserts as well as their respective culture medium were provided by Epithelix Sàrl (Geneva, Switzerland). HAEs were cultured in 24-well inserts at the air-liquid interface. All cells and HAEs were maintained at 37°C with 5% CO_2_.

Pandemic influenza A/California/7/2009 (A(H1N1)pdm09) virus was amplified on MDCK α2-6 cells in MEM supplemented with 1% HEPES and 1 μg/mL trypsin treated with N-tosyl-L-phenylalanine chloromethyl ketone (Sigma-Aldrich, Oakville, ON, Canada). For both *in vitro* and *in vivo* experiments, culture medium of cells infected with IAV was collected and centrifuged to remove cell debris. Viral titers were determined by plaque assays. The RSV-A2 strain (VR-1540; ATCC) was amplified in HEp-2 cells in MEM supplemented with 2% FBS and 1% HEPES. For experiments with HAEs, culture medium of cells infected with RSV was collected and centrifuged to remove cell debris. For animal experiments, clarified supernatant was ultracentrifuged and the pellet was resuspended in fresh culture medium to get a sufficiently high viral inoculum. Viral titers were determined by immunostaining with a goat anti-RSV primary antibody (B65860G; Meridian; Cedarlane Laboratories Ltd., Burlington, ON, Canada) and a horseradish peroxidase-labeled rabbit anti-goat IgG secondary antibody (HAF017; R&D Systems; Cedarlane Laboratories Ltd.) as described (56). Infectious foci were revealed by using the True-Blue™ Peroxidase Substrate (KPL, Gaithersburg, MD, USA).

### Infection kinetics in mice

Our study exclusively examined female mice. It is unknown whether the findings are relevant for male mice. Six week-old female BALB/c mice (Charles River, Saint-Constant, QC, Canada) were anesthetized with isoflurane and infected with 1.0×10^6^ plaque forming units (PFUs) of RSV-A2, 5.0×10^4^ PFUs of A(H1N1)pdm09 (non-lethal doses that result in a significant weight loss and a significant lung viral load), or simultaneously with both viruses at the same inoculum in a total volume of 50 µL of MEM that was delivered by instillation in both nostrils. Mock-infected mice received a total volume of 50 µL of MEM intranasally. For sequential infections, mice were infected with both viruses either 1 day or 4 days apart. Groups of mice also received a first intranasal administration of MEM, RSV-A2 or A(H1N1)pdm09 followed by MEM 1 day or 4 days apart.

Animals were monitored for clinical signs of infection (weight loss, labored breathing, reduced grooming, and hunched posture) daily for 14 days. Mice were euthanized when the weight loss was greater or equal to 20% of initial weight or when two other sickness signs reached a limit point. On day 4 after the last infection, lungs were collected, frozen in liquid nitrogen and stored at -80°C for quantification of viral RNA load, IFN and ISG mRNAs as well as for cytokine protein levels. Some lungs were also injected with 4% paraformaldehyde, collected, and stored in 4% paraformaldehyde at 4°C for histopathological evaluation.

On days 1, 4 and 7 p.i., lungs of mice infected with RSV and IAV were collected, weighed (wet lung), and dried for 24 h before weighing again (dry lung). The lung wet-to-dry weight ratio was then calculated and compared with that of non-infected mice.

All animals were used in accordance with the Canadian Council on Animal Care guidelines, and the protocol was approved by the Animal Care Ethics Committee of Laval University (protocol 2022-1159).

### Tissue homogenization and RNA isolation

Lungs were homogenized in 1 mL sterile phosphate-buffered saline (PBS) containing gentamicin (2 mg/mL; Life Technologies, Burlington, ON, Canada), vancomycin (25 mg/mL; Sterimax Inc., Oakville, ON, Canada) and fungizone (0.125 mg/mL; Wisent Inc., Saint-Jean-Baptiste, QC, Canada) on ice using a tissue homogenizer (OMNI International, Ottawa, ON, Canada). Homogenates were centrifuged (2 500 rpm) for 10 min at 4°C. The supernatant was then collected and divided in separate samples before being stored at -80°C. A sample of 200 µL was treated with protease and phosphatase inhibitors by adding 20 µL of 10× solution of PhosStop EASYpack and cOmplete Mini EDTA-free EASYpack (both from Roche Diagnostics, Laval, QC, Canada) diluted in PBS for cytokine quantification by immunoassay. Another 100 µL was transferred in a tube containing 850 µL of Trizol reagent (Life Technologies) and RNA isolation was performed using the Direct-zol™ RNA MiniPrep Plus kit (Zymo Research; Cedarlane Laboratories Ltd.) according to the manufacturer’s instructions for quantification of viral RNA load and IFN-α/β/λ2/3 mRNAs.

### Infection kinetics in HAEs

Prior to infection, the apical pole of HAEs was incubated with 200 µL of pre-warmed Opti-MEM (Gibco; ThermoFisher Scientific, Waltham, MA, USA) for 10 min at 37°C to remove the mucus layer. The apical wash was then taken after pipetting up and down a few times. HAEs were infected at the apical pole with RSV-A2 or A(H1N1)pdm09 at a multiplicity of infection (MOI) of 0.02 (considering that each HAE was made of 500 000 cells), in a volume of 200 µL of Opti-MEM. HAEs were incubated for 1 h at 37°C with 5% CO_2_ and, the viral suspension was then removed. For simultaneous coinfections, both viruses at the same MOI (0.02) were added in a total volume of 200 µL of medium. Sequential infections with both viruses were performed 1 day or 4 days apart. Fig. 2 summarizes the experimental design.

Each day following the first infection, the apical pole of HAEs was incubated with 200 µL of pre-warmed Opti-MEM for 10 min at 37°C with 5% CO_2_. The apical wash was then collected after pipetting up and down a few times, and stored at -80°C until use. Total RNA was extracted from 100 µL of HAE apical wash using the EZ2 Connect system (EZ1&2 Virus Mini Kit v2.0, Qiagen, Toronto, ON, Canada) for viral RNA load quantification. Every 48 h, the basolateral medium was collected and replaced with 500 µL of fresh pre-warmed culture medium. The collected basolateral medium was snap-frozen and stored at -80°C for cytokine protein quantification. At selected time points, 100 µL of AVL lysis buffer (Qiagen) were added onto the apical pole of HAEs for 30 min at room temperature to lyse the cells. Total RNA was extracted from cell lysates using the EZ2 Connect system (EZ1&2 Virus Mini Kit v2.0, Qiagen) for ISG and 18S mRNA quantification by RT-ddPCR. Mock-infected HAEs were processed as infected HAEs for 5 days before lysis.

### Treatment of HAEs with recombinant IFN proteins

Recombinant human IFN-α2a (H6041; Sigma-Aldrich) was reconstituted in PBS with 0.1% bovine serum albumin (BSA; Sigma-Aldrich) then added to the basolateral pole at a concentration of 100 U/mL. Recombinant human IFN-β (8499-IF) and IFN-λ2 (1587-IL) were purchased from R&D Systems (Minneapolis, MN, USA) and were reconstituted in PBS with 0.1% BSA. They were added to the basolateral pole at a concentration of 100 ng/mL. Nasal HAEs were treated with recombinant proteins 1 day prior to their infection with RSV-A2 or A(H1N1)pdm09 at a MOI of 0.02 and then daily until 5 days p.i.

### Viral RNA load by RT-qPCR

Viral load was quantified in total RNA extracted (5 µL) from either mouse lung homogenates or HAE apical washes by RT-qPCR assays performed with the QuantiTect Virus + ROX Vial Kit (Qiagen) in a LightCycler^®^ 480 system (Roche Molecular System, Laval, QC, Canada). Primers and probes targeting the nucleocapsid protein (N) gene of RSV-A2 (57) and the matrix protein (M) gene of influenza A (sequences available upon request) were used.

### Expression of murine IFN-**α**/**β**/**λ** and human ISGs by RT-ddPCR

For the quantification of IFN-α/β/λ2/3 mRNAs (from mouse lung homogenates) as well as the 18S housekeeping gene mRNAs (from mouse lung homogenates and HAE cell lysates), reverse transcription was performed using the SuperScript™ IV First-Strand Synthesis System (Invitrogen™, ThermoFisher Scientific) according to the manufacturer’s instructions using 5 µL of extracted RNA. Droplet digital PCR reactions were performed using QX200™ ddPCR™ EvaGreen SuperMix (Bio-Rad Laboratories Ltd., Mississauga, ON, Canada). Primers are described in Table S1.

Expression of ISGs in lysates of nasal HAEs were determined with the One-step RT-ddPCR Advanced Kit for probes (Bio-Rad Laboratories Ltd.), using primers and probes targeting 2’,5’-oligoadenylate synthetase 1 (OAS1), interferon-induced transmembrane protein 3 (IFITM3), interferon-stimulated gene 15 (ISG15), and myxovirus resistance protein A (MxA) using 5 µL of extracted RNA. Primers and probes are described in S1 Table.

For all ddPCR reactions, droplets were generated using a QX200™ Droplet Generator (Bio-Rad Laboratories Ltd.) and PCR reactions were performed using a C1000 Touch Thermal cycler (Bio-Rad Laboratories Ltd.). Acquisition was made with a QX200™ Droplet Reader (Bio-Rad Laboratories Ltd.), with the software QX Manager 1.2.

### Cytokine protein quantification by magnetic bead-based immunoassay

Cytokine proteins were quantified in mouse lung homogenates clarified by centrifugation (100 µL) with a Bio-Plex™ Pro Mouse Cytokine Express assay for six targets (IFN-γ, IL-1α, IL-1β, IL-6, IL-10 and TNF-α) (Bio-Rad Laboratories Ltd.) according to the manufacturer’s instructions.

IFN proteins were quantified in 250 µL of HAE basolateral medium samples with a Bio-Plex™ Pro Human Inflammation Panel 1 Express assay for three targets (IFN-β, IL-28A/IFN-λ2 and IL-29/IFN-λ1) (Bio-Rad Laboratories Ltd.) according to the manufacturer’s instructions.

A Bioplex 200 system and the Bioplex Manager Software V6.2 (Bio-Rad Laboratories Ltd.) were used to measure the mean fluorescence intensity from all the bead combinations.

### Histopathology

Formalin-fixed paraffin-embedded lung tissues were cut in 4 µm-thick histologic sections and stained with hematoxylin-eosin. For microphotographs, slides were digitalized at 40× magnification using a Nanozoomer slide scanner (Hamamatsu, Shizuoka, Japan) and NDP viewer 2.0 software (Hamamatsu, Japan). Scoring of histologic parameters was performed by a pathologist (CC), blind to experimental groups. A semiquantitative scale (0=normal/absent; 1=mild; 2=moderate and 3=marked) was used to score the intensity of bronchial/endobronchial, peribronchial, perivascular, interstitial, pleural and intra-alveolar inflammation (58).

## Statistical analyses

Statistical analyses were performed with GraphPad Prism version 9.4.0 (GraphPad Software, La Jolla, CA, USA). A Shapiro-Wilk normality test was first performed on all data. Percentage of body weight changes were compared using a two-way analysis of variance (ANOVA) with a Tukey’s multiple comparison test. For all other data, normal data were analyzed using a one-way Brown-Forsythe and Welch ANOVA test with post-hoc Dunnett’s T3 multiple comparison test or a Welch’s t-test when comparing only two conditions. For data that failed the normality test or when the N was too small for normality assessment, a non-parametric Kruskal-Wallis ANOVA with a Dunn’s multiple comparison test was used.

## Data availability statement

Data will be made available upon reasonable request to the corresponding author.

## Conflict of interest statement

The authors have declared that no conflict of interest exists.

## Acknowledgments

We acknowledge the bioimaging platform of the Infectious Disease Research Centre, funded by an equipment and infrastructure grant from the Canadian Foundation of Innovation.

This work was supported by a Foundation Grant from the Canadian Institutes of Health Research [grant no. 148361 to G.B.].

## Notes

### Competing Interest Statement

The authors have declared no competing interest.

