## Supplementary material for "Influenza A Virus Interferes With Respiratory Syncytial Virus in Mice and Reconstituted Human Airway Epithelium"

**Supplemental material:**

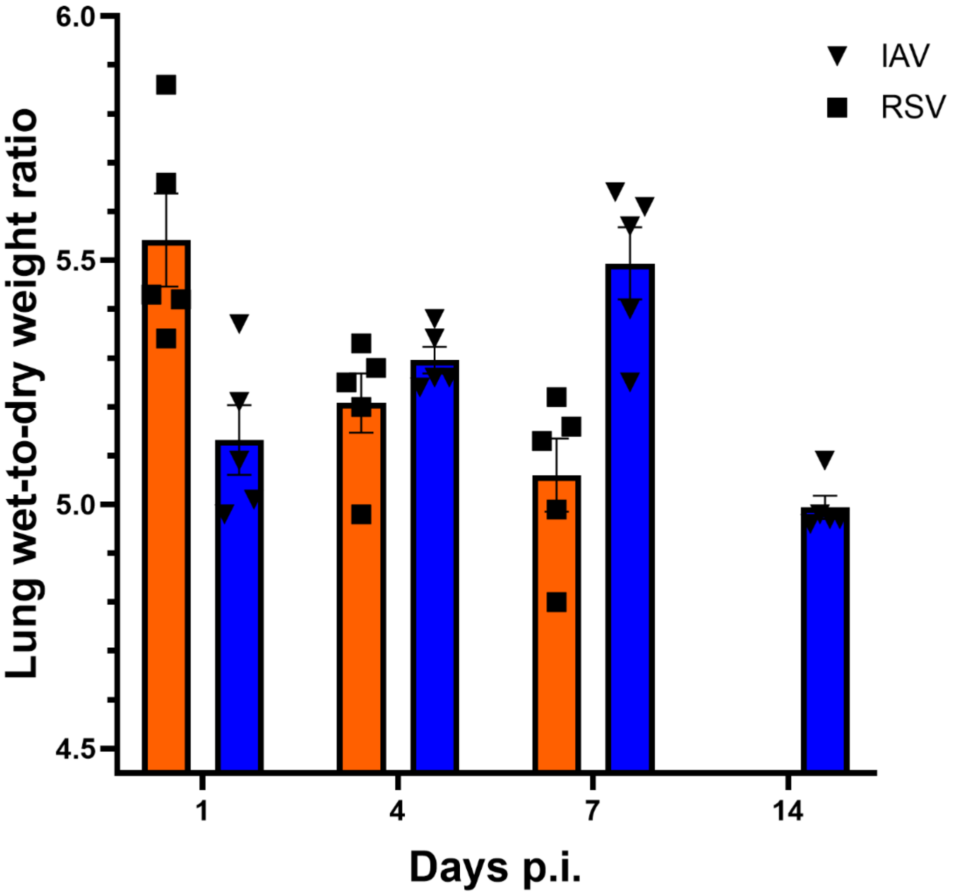


**FIG S1. Lung wet-to-dry weight ratio in mice infected with respiratory syncytial virus (RSV) or influenza virus (IAV).** Mice were infected intranasally with RSV or IAV. On days 1, 4, 7 and 14 (IAV only) post-infection, lungs were collected and weighed (wet lung). Lungs were dried for 24 h and weighed again (dry lung) and the wet-to-dry weight ratio was calculated. The ordinate shows the increase in wet-to-dry weight ratio over non-infected controls (4.45±0.46). Results are expressed as the mean ± SEM of 5 mice per group from a single experiment.

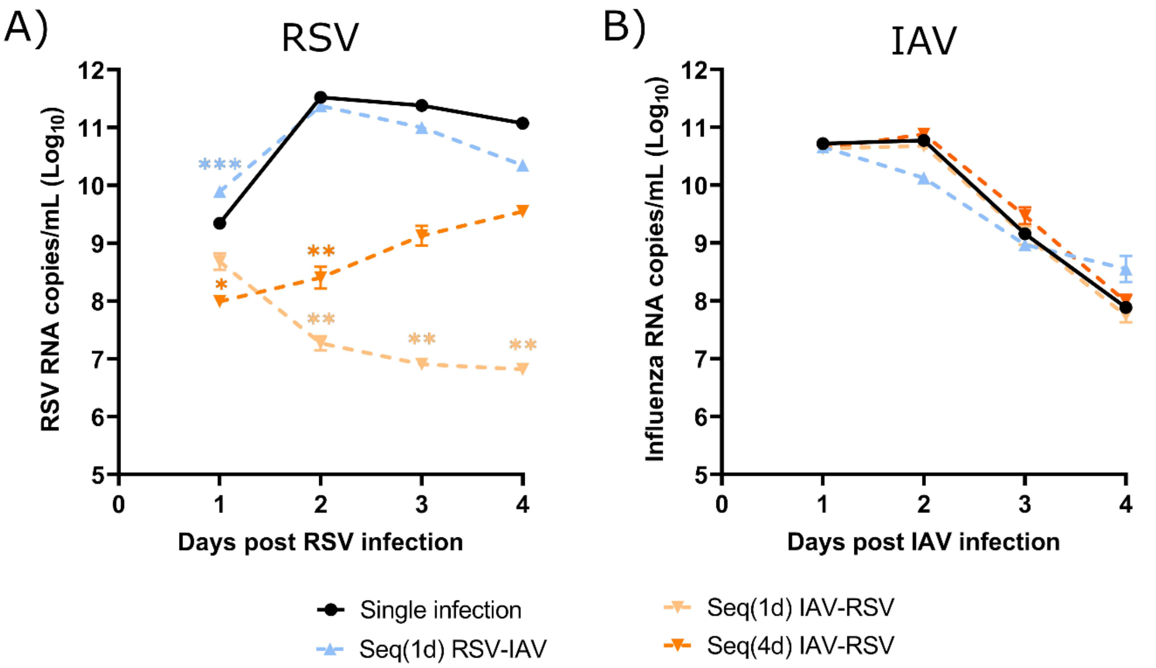


**FIG S2. Viral interference between respiratory syncytial virus (RSV) and influenza A virus (IAV) in bronchiolar human airway epithelia (HAEs).** Bronchiolar HAEs were infected with RSV, IAV, or both viruses simultaneously or sequentially with a 1-day (Seq(1d)) or a 4-day (Seq(4d)) interval. Viral RNA loads were determined in apical washes by RT-qPCR on a daily basis. Days post-infection (p.i.) represent the time after infection with either RSV (A) or IAV (B). Results are expressed as the mean of the $Log_{10}$ of viral RNA copies per mL ± SEM of 3-4 bronchiolar HAE inserts from two independent experiments. *, $p \leq 0.05$; **, $p \leq 0.01$; ***, $p \leq 0.001$ compared to single infection.

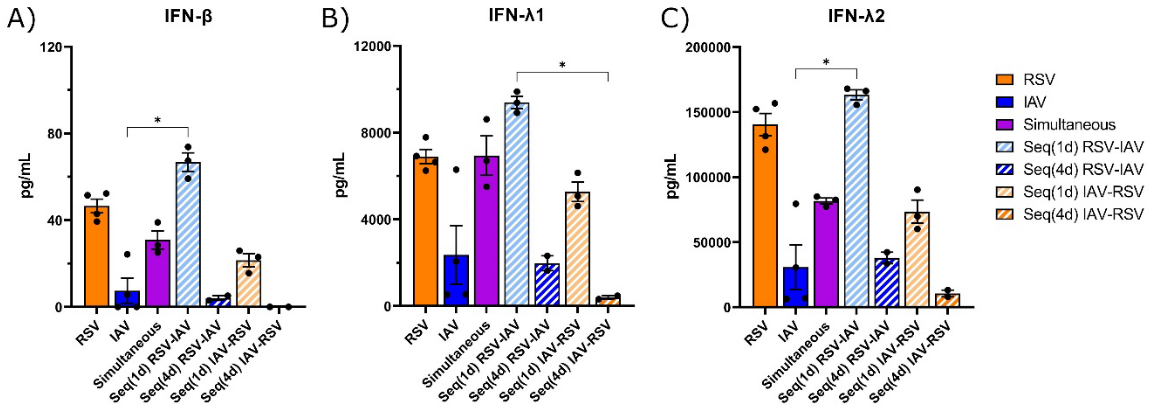


**FIG S3. Interferon (IFN) protein production in nasal human airway epithelia (HAEs) infected with respiratory syncytial virus (RSV), influenza A virus (IAV) or both viruses.** Nasal HAEs were infected with RSV, IAV, or both viruses simultaneously or sequentially with a 1-day (Seq(1d)) or a 4-day (Seq(4d)) interval. IFN-β (A), IFN-λ1 (B) and IFN-λ2 (C) protein production was measured in basolateral medium by magnetic microbead immunoassays on day 5 after the last infection, except for the Seq(1d) groups (4 days after the second viral challenge). Results are expressed as the mean amount of IFN proteins in pg per mL ± SEM of 2-4 nasal HAE inserts from two independent experiments. *, $p \leq 0.05$ between indicated groups.

**Table S1. Sequences of primers and probes used for the quantification of murine interferons, human interferon-stimulated genes and 18S housekeeping gene by RT-ddPCR.**

<table>
<tr><th></th><th></th><th>Sequence</th><th>Reference</th></tr>
<tr><td rowspan="3">IFN-α</td><td>Forward 1</td><td>5'-CTG ACC CAG GAA GAC TCC CTG-3'</td><td>Enlow et al., 2021[a]</td></tr>
<tr><td>Forward 2</td><td>5'-CTG ACC CAG GAA GAC GCC CTG-3'</td><td>Enlow et al., 2021[a]</td></tr>
<tr><td>Reverse</td><td>5'-GAC TTC TGC TCT GAC CAC CTC-3'</td><td>Enlow et al., 2021a</td></tr>
<tr><td rowspan="2">IFN-β</td><td>Forward</td><td>5'-CTA TTG TTG TAC GTC TCC TGG-3'</td><td>Enlow et al., 2021[a]</td></tr>
<tr><td>Reverse</td><td>5'-GGC GTA GCT GTT GTA CTT CAT GA-3'</td><td>Enlow et al., 2021[a]</td></tr>
<tr><td rowspan="2">IFN-λ2/3</td><td>Forward</td><td>5'-CTGMTGGCCGCAGTGCTGA-3'</td><td>This work</td></tr>
<tr><td>Reverse</td><td>5'-CAGCTGCYTCAGGTCCCAG-3'</td><td>This work</td></tr>
<tr><td rowspan="3">OAS1</td><td>Forward</td><td>5'-GCA AAC AGG TCT GGG AGG-3'</td><td>Gilbert-Girard et al., 2024[b]</td></tr>
<tr><td>Reverse</td><td>5'-GTC AAT GGC ATG GTT GAT TTG C-3'</td><td>Gilbert-Girard et al., 2024[b]</td></tr>
<tr><td>Probe</td><td>5'-CAG TTC TGT TGC CAC TCT CTC TCC TG-3'</td><td>Gilbert-Girard et al., 2024[b]</td></tr>
<tr><td rowspan="3">IFITM3</td><td>Forward</td><td>5'-ATC GTC ATC CCA GTG CTG AT-3'</td><td>Cheemarla et al., 2021[c]</td></tr>
<tr><td>Reverse</td><td>5'-ATG GAA GTT GGA GTA CGT GG-3'</td><td>Cheemarla et al., 2021[c]</td></tr>
<tr><td>Probe</td><td>5'-CAG GAG GCA TCA CTG AGG CCA G-3'</td><td>Gilbert-Girard et al., 2024[b]</td></tr>
<tr><td rowspan="3">ISG15</td><td>Forward</td><td>5'-TGG ACA AAT GCG ACG AAC C-3'</td><td>Gilbert-Girard et al., 2024[b]</td></tr>
<tr><td>Reverse</td><td>5'-GGT CAG CCA GAA CAG GTC-3'</td><td>Gilbert-Girard et al., 2024[b]</td></tr>
<tr><td>Probe</td><td>5'-CTG GTG AGG AAT AAC AAG GGC CGC-3'</td><td>Gilbert-Girard et al., 2024[b]</td></tr>
<tr><td>MxA</td><td>Forward</td><td>5'-GTC AGT TAC CAG GAC TAC GA-3'</td><td>Gilbert-Girard et al., 2024[b]</td></tr>
</table>

| | | | |
|---|---|---|---|
| | Reverse | 5'- ATC TCC AGG GTG ATT AGC TC-3' | Gilbert-Girard et al., 2024[b] |
| | Probe | 5'-TTG AGA TTT CGG ATG CTT CAG AGG TAG-3' | Gilbert-Girard et al., 2024[b] |
| **18S** | Forward | 5'-GGA TGC GTG CAT TTA TCA G-3' | Gilbert-Girard et al., 2024[b] |
| | Reverse | 5'-AGT TGA TAG GGC AGA CGT TC-3' | Gilbert-Girard et al., 2024[b] |

IFITM3, interferon-induced transmembrane protein 3; IFN, interferon; ISG15, interferon-stimulated gene 15; MxA, myxovirus resistance protein A; OAS1, 2',5'-oligoadenylate synthetase 1.

[a], Enlow W, Bordeleau M, Piret J, Ibáñez FG, Uyar O, Venable MC, Goyette N, Carbonneau J, Tremblay ME, Boivin G. Microglia are involved in phagocytosis and extracellular digestion during Zika virus encephalitis in young adult immunodeficient mice. J Neuroinflammation. 2021; 18:178. doi: 10.1186/s12974-021-02221-z. PubMed PMID 3439977.

[b], Gilbert-Girard S, Piret J, Carbonneau J, Henaut M, Goyette N, Boivin G. 2024. Viral interference between severe acute respiratory syndrome coronavirus 2 and influenza A viruses. PLoS Pathog 20:e1012017.

[c], Cheemarla NR, Watkins TA, Mihaylova VT, Wang B, Zhao D, Wang G, et al. Dynamic innate immune response determines susceptibility to SARS-CoV-2 infection and early replication kinetics. J Exp Med. 2021; 218:e20210583. Epub 20210615. doi: 10.1084/jem.20210583. PubMed PMID: 34128960.
